# “ER-Ca^2+^ sensor STIM regulates neuropeptides required for development under nutrient restriction in *Drosophila*”

**DOI:** 10.1101/386649

**Authors:** Megha, Christian Wegener, Gaiti Hasan

## Abstract

Neuroendocrine cells communicate via neuropeptides to regulate behaviour and physiology. This study examines how STIM (Stromal Interacting Molecule), an ER-Ca^2+^ sensor required for Store-operated Ca^2+^ entry, regulates neuropeptides required for *Drosophila* development under nutrient restriction (NR). We find two STIM-regulated peptides, Corazonin and short Neuropeptide F, to be required for NR larvae to complete development. Further, a set of secretory DLP (Dorso lateral peptidergic) neurons which co-express both peptides was identified. Partial loss of *dSTIM* caused peptide accumulation in the DLPs, and reduced systemic Corazonin signalling. Upon NR, larval development correlated with increased peptide levels in the DLPs, which failed to occur when *dSTIM* was reduced. Comparison of systemic and cellular phenotypes associated with reduced *dSTIM*, with other cellular perturbations, along with genetic rescue experiments, suggested that *dSTIM* primarily compromises neuroendocrine function by interfering with neuropeptide release. Under chronic stimulation, *dSTIM* also appears to regulate neuropeptide synthesis.

## Introduction

Metazoan cells commonly use ionic Ca^2+^ as a second messenger in signal transduction pathways. To do so, levels of cytosolic Ca^2+^ are dynamically managed. In the resting state, cytosolic Ca^2+^ concentration is kept low and maintained thus by the active sequestration of Ca^2+^ into various organelles, the largest of which is the ER. Upon activation, ligand-activated Ca^2+^ channels on the ER, such asthe ryanodine receptor or inositol 1,4,5-trisphosphate receptor (IP_3_R), release ER-store Ca^2+^ into the cytosol. Loss of ER-Ca^2+^ causes STromal Interacting Molecule (STIM), an ER-resident transmembrane protein, to dimerize and undergo structural rearrangements. This facilitates the binding of STIM to Orai, a Ca^2+^ channel on the plasma membrane, whose pore now opens to allow Ca^2+^ from the extracellular milieu to flow into the cytosol. This type of capacitative Ca^2+^ entry is called Store-operated Ca^2+^ entry (SOCE) [1]. Of note, key components of SOCE include the IP_3_R, STIM and Orai, that are ubiquitously expressed in the animal kingdom, underscoring the importance of SOCE to cellular functioning. Depending on cell type and context, SOCE can regulate an array of cellular processes [2].

Neuronal function in particular is fundamentally reliant on the elevation of cytosolic Ca^2+^. By tuning the frequency and amplitude of cytosolic Ca^2+^ signals that are generated, distinct stimuli can make the same neuron produce outcomes of different strengths [3]. The source of the Ca^2+^ influx itself contributes to such modulation as it can either be from internal ER-stores or from the external milieu, through various activity-dependent voltage gated Ca^2+^ channels (VGCCs) and receptor-activated Ca^2+^ channels ora combination of the two. Although the contributions of internal ER-Ca^2+^ stores to neuronal Ca^2+^ dynamics are well recognized, the study of how STIM and subsequently, SOCE-mediated by it, influences neuronal functioning, is as yet a nascent field.

Mammals have two isoforms of STIM, STIM1 and STIM2, both which are widely expressed in the brain. As mammalian neurons also express multiple isoforms of Orai and IP_3_R, it follows that STIM-mediated SOCE might occur in them. Support for this comes from studies in mice, where STIM1-mediated SOCE has been reported for cerebellar granule neurons [4] and isolated Purkinje neurons [5], while STIM2-mediated SOCE has been shown in cortical [6] and hippocampal neurons [7]. STIM can also have SOCE-independent roles in excitable cells, that are in contrast to its role via SOCE. In rat cortical neurons [8] and vascular smooth muscle cells [9], Ca^2+^ release from ER-stores prompts the translocation of STIM1 to ER-plasma membrane junctions, and binding to the L-type VGCC, Ca_v_1.2. Here STIM1 inhibits Ca_v_1.2 directly and causes it to be internalized, reducing the long-term excitability of these cells. In cardiomyocyte-derived HL1 cells, STIM1 binds to a T-type VGCC, Ca_v_1.3, to manage Ca^2+^ oscillations during contractions [10]. These studies indicate that STIM regulates cytosolic Ca^2+^ dynamics in excitable cells, including neurons and that an array of other proteins determines if STIM regulation results in activation or inhibition of neurons. Despite knowledge of the expression of STIM1 and STIM2 in the hypothalamus (Human Protein Atlas), the major neuroendocrine centre in vertebrates, studies on STIM in neuroendocrine cells are scarce. We therefore used *Drosophila melanogaster (Drosophila)*, the vinegar fly, to address this gap.

Neuroendocrine cells possess elaborate machinery for the production, processing and secretion of neuropeptides (NPs), which perhaps form the largest group of evolutionarily conserved signalling agents [11,12]. Inside the brain, NPs typically modulate neuronal activity and consequently, circuits; when released systemically, they act as hormones. *Drosophila* is typical in having a vast repertoire of NPs that together play a role in almost every aspect of its behaviour and physiology [13,14]. Consequently, NP synthesis and release are highly regulated processes. As elevation in cytosolic Ca^2+^ is required for NP release, a contribution for STIM-mediated SOCE to NE function was hypothesized.

*Drosophila* possess a single gene for STIM, IP_3_R and Orai, and all three interact to regulate SOCE in *Drosophila* neurons [15,16]. In dopaminergic neurons, *dSTIM* is important for flight circuit maturation [15–17], with dSTIM-mediated SOCE regulating expression of a number of genes, including *Ral*, which controls neuronal vesicle exocytosis [17]. In glutamatergic neurons, *dSTIM* is required for development under nutritional stress and its’ loss results in down-regulation of several ion channel genes which ultimately control neuronal excitability [18]. Further, *dSTIM* over-expression in insulin-producing NE neurons could restore Ca^2+^ homeostasis in a non-autonomous manner in other neurons of an IP_3_R mutant [19], indicating an important role for dSTIM in NE cell output, as well as compensatory interplay between IP_3_R and dSTIM. At a cellular level, partial loss of dSTIM impairs SOCE in *Drosophila* neurons [15,17,20] as well as mammalian neural precursor cells [21]. Additionally, reducing dSTIM in *Drosophila* dopaminergic neurons attenuates KCI-evoked depolarisation and as well as vesicle release [17]. Because loss of dSTIM specifically in *dimm^+^* NE cells results in a pupariation defect on nutrient restricted (NR) media [22], we used the NR paradigm as a physiologically relevant context in which to investigate STIM’s role in NE cells from the cellular as well as systemic perspective.

## Results

### SOCE is required in sNPF and Crz producing cells for development under nutritional stress

Collectively, more than 20 different NPs are known to be made by the neuroendocrine cells in which reducing SOCE components resulted in poor pupariation upon NR [22]. To shortlist specific NPs important for this paradigm, we undertook a curated *GAL4*-*UAS* screen. NP-GAL4S were used to drive the knockdown of *IP*_*3*_*R* (*IP*_*3*_*R*^*IR*^) [23], and pupariation of the resulting larvae were scored on normal vs NR media (Fig. S1A). On normal food, a significant reduction of pupariation was seen only with *sNPF-GAL4*(Fig. S1A), whose expression strongly correlates with neurons producing sNPF [24]. Upon NR, the largest effect was seen with *sNPF-GAL4*, followed by small but significant pupariation defect with *AstA-GAL4* and *DSK-GAL4* (Fig. S1A). Neurons that secrete NPs may also secrete neurotransmitters, therefore, a role specifically for sNPF was tested. Reducing the level of *sNPF*(*sNPF*^*IR*^) or reducing an enzyme required for neuropeptide processing (*amontillado*; *amon*^*IR*^) [25] in *sNPF-GAL4* expressing cells, as well as a hypomorphic sNPF mutation (*sNPF*^*00448*^) resulted in impairment of larval development upon NR (Fig. S1B). These data indicate that sNPF is required for pupariating under NR conditions.

*sNPF-GAL4* expresses in large number of neurons (>300) in the larval brain [24] (Fig. S1C), and also expresses in the larval midgut and epidermis. To further refine sNPF^+^ neurons on which we can perform cellular investigations, we tested a *Crz-GAL4* driver. This driver expresses in fewer neurons (~22), all of which express the neuropeptide Corazonin (Crz). Importantly, a small subset of these, three bilateral neurons in the brain lobe, make Crz and sNPF. [24] (Fig. 1A). Reducing SOCE in Crz neurons, by reducing either IP_3_R or STIM (*dSTIM*^*IR*^) [15,16] or over-expressing a dominant-negative version of Orai (*Orai*^*E18oA*^) [26], resulted in reduced pupariation on NR (Fig. 1B). The absence of a developmental defect on normal food suggests that SOCE in these neurons is primarily required to survive NR.

To test if both sNPF and Crz were required, they were specifically reduced (*sNPF*^*IR*^; *Crz*^*IR*^) in Crz neurons. Knockdown of either NP resulted in larvae with a pupariation defect on NR media but not, on normal food (Fig. 1C; Fig. S1D). In *Drosophila* neurons, enhancing the expression of SOCE regulators leads to increased SOCE [16]. To test the positive effect of SOCE on Crz and sNPF, a genetic compensation experiment was carried out. The SOCE-regulators, *IP*_*3*_*R* or *dSTIM* were over-expressed in Crz neurons which also expressed reduced levels of either sNPF or Crz. NR larvae with this genetic make-up showed a significant improvement in pupariation on NR media, as compared to NR larvae with only reduced NPs (Fig. 1C). Interestingly, the compensation was sufficient to also increase the number of adults that emerged (Fig. 1D). Notably, over-expression of either of the two SOCE molecules, dSTIM and IP_3_R on their own, did not affect pupariation on either normal or NR media (Fig. 1C), but unlike on normal food (Fig. S1E), did reduce development to adulthood on NR media (Fig. 1D). These data underscore the sensitivity of Crz neurons to ER-Ca^2+^ homeostasis during NR.

Loss of IP_3_R and sNPF has previously been shown to affect larval feeding [27–29]. Hence, larval intake of dye-colored food in a 2-hour span was measured. Age-synchronized larvae with knockdown of either *dSTIM*, *IP*_*3*_*R*, *Crz* or *sNPF* in Crz neurons exhibited no difference in the amount of dye ingested (Fig. S1F), suggesting that developmental defects in the NR assay do not arise from a fundamental feeding problem.

Altogether, these genetic experiments helped identify a set of NP expressing neurons, the Crz neurons, a subset of which also express sNPF, in which SOCE plays an important role during development on NR.

**Fig. 1.**
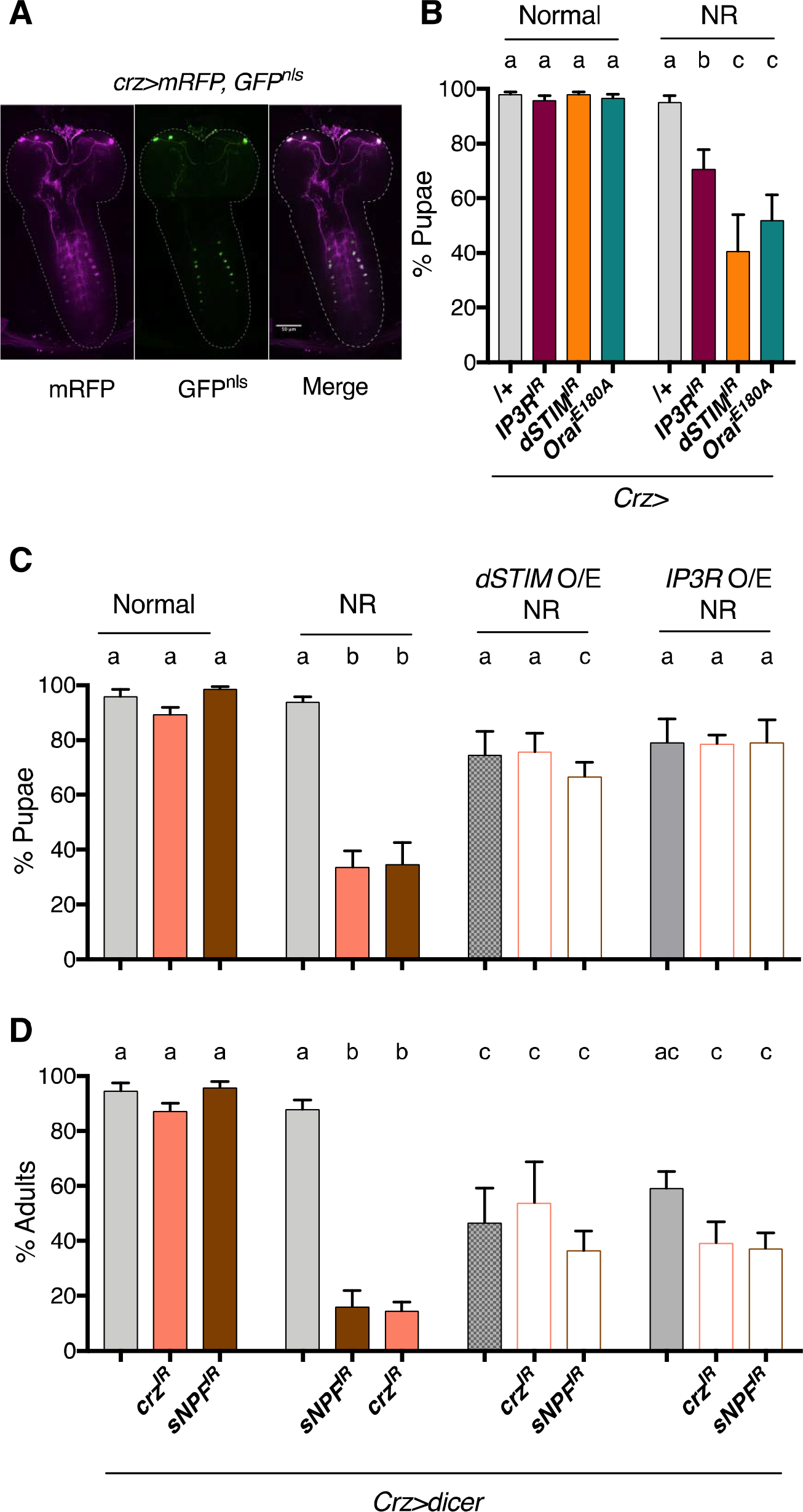
SOCE is required in Crz neurons for larval development on NR media. **(A)** Expression pattern of *Crz-GAL4* driver, used in this study to manipulate Crz neurons, visualised by expressing membrane bound RFP (mRFP) and GFP with a nuclear localisation signal (GFP^nls^) **(B)** % Pupae upon reduction of SOCE by knockdown of STIM (*STIM*^*IR*^), *IP*_*3*_*R* (*IP*_*3*_*R*^*IR*^) or ectopic expression of a dominant-negative *Orai* (*Orai*^*E18oA*^) in Crz neurons. To measure pupariation, twenty five, 88h±3hold larvae, per vial, were transferred to either normal food (corn flour, yeast, sugar) (See materials and methods for exact composition) or nutrient restricted (NR; 100mM Sucrose) media and number of pupae (and adults where relevant) that developed were counted. N = 6 vials for all experiments in this study. **(C)** % Pupae upon reduction of either *sNPF* or *Crz* (*Crz*^*IR*^, *sNPF*^*IR*^) in Crz neurons, and when, dSTIM or *IP*_*3*_*R* are expressed in this background (*dSTIM* O/E; *IP*_*3*_*R* O/E). **(D)** % Adults recovered for genotypes in **(C)**. Bars with the same alphabet represent statistically indistinguishable groups. Two-way ANOVA with Sidak multi comparison p<0.05 for (B), (C) and (D). See also Figure Supplement 1.

### Crz^+^ and sNPF^+^ DLP neurons majorly contribute to development on NR, and are activated by NR

In the larval CNS, Crz is expressed in 3 pairs of DLPs (Dorso Lateral Peptidergic neurons) in the *pars lateralis* region of the brain lobes, 1 pair of neurons in dorso-medial region and 8 pairs of interneurons in the VG (ventral ganglion) [30]. Other than the dorso-medial neurons, the *Crz-GAL4* used in this study recapitulates the known expression pattern for Crz. (Fig. S2A, B; Cartoon: Fig. 2A). Additionally, adjacent to DLPs, low expression of *Crz-GAL4* was observed in 3-4 neurons that do not express Crz (Fig. S2A and Fig. S2C). As mentioned previously, Crz^+^ DLPs co-express sNPF [24]. In terms of neuronal architecture, the DLP neurons have two major branches: the anterior branch culminates in a dense nest of neurites at the ring gland (RG), while the posterior branch terminates in the subesophageal zone (SEZ). The VG neurons form a network amongst themselves to ultimately give rise to two parallel bundles that travel anteriorly, and end in the brain lobes. To visualize the overall distribution of NPs in the Crz neurons, we ectopically expressed a rat neuropeptide coupled to GFP (ANF∷GFP), a popular tool used to track NP transport and release in *Drosophila* [31] (Fig. S2D). Firstly, within the DLPs, like Crz∷mcherry (Fig. S2A), ANF∷GPF was either in the cell bodies or RG projections, but not in the projections terminating at the subesophageal zone (Fig. S2D), suggesting selective NP transport to the RG, which is a major neurohaemal site for systemic release of neuropeptides. Secondly, ANF∷GFP intensity was higher in the cell bodies of the DLPs than VG neurons (Fig. S2D).

The close proximity of the terminal projections of the Crz^+^ VG neurons and the anterior branch of the Crz^+^ DLP neurons in the brain lobe suggested possible neuromodulation between the two sets of neurons. Therefore, we undertook experiments to distinguish the contribution of DLPs vs VG localized Crz neurons, to the development in NR media. First, we utilized *tshGAL80* to restrict *Crz-GAL4* expression to the DLPs (Fig. S2E). The level of pupariation under NR conditions observed with restricted expression of *dSTIM*^*IR*^ (Fig. 2B; Mean: 51%±4.2) was similar to that seen with full expression (Fig. 1B; Mean: 40.7%±13.3), suggesting a major contribution of the DLP neurons to the NR phenotype. Furthermore, *sNPF*>*Crz*^*IR*^ larvae have levels of pupariation of NR larvae (Fig. S2F; Mean: 30.9%±7.8) similar to *Crz*>*Crz*^*IR*^ NR larvae (Fig. 1C; Mean: 33.8%±5.9). Because *sNPF-GAL4* marks only the Crz^+^ DLP neurons and not the Crz^+^ VG neurons (Fig. S1C), this too suggests a major role for the Crz^+^ DLP neurons.

Requirement of SOCE in Crz neurons for pupariation on NR (Fig. 1C) suggested that these neurons experience elevated cytosolic Ca^2+^ in NR conditions and are therefore, stimulated by chronic starvation. To test this, the UV light-activated genetically encoded calcium sensor, CaMPARI [32], was utilised. The sensor fluoresces in the GFP range (F_488_) and is converted irreversibly to fluoresce in the RFP range (F_561_), when exposed to UV light and in the presence of Ca^2+^. The level of conversion positively titrates with Ca^2+^ concentrations. Larva expressing CaMPARI in Crz^+^ neurons were placed in either normal or NR media for 24 hours (24h NR). Whole larvae were immobilized, and exposed to UV light for 2mins. Control larva were subject to the same treatment but, without being exposed to UV light (Fig. 2C,D; no UV). Detection of F_561_ in the fed state suggests that these neurons are active even under normal food conditions (Fig. 2C,D). Notably, after 24 hours on NR media, chronic starvation caused a ~2-fold increase in average levels F_561_ and therefore, of neuronal activation (Fig. 2D). F_561_/F_488_ ratios did not appear to change in the VG neurons (Fig. S2G,H). While there is a possibility that VG neurons do not exhibit higher F_561_ because of insufficient penetration of UV light, the CaMPARI results together with the genetic experiments (Fig. 2B, S2F), formed the basis for selecting the DLP neurons for further analysis on how dSTIM affects Crz and sNPF.

**Fig. 2.**
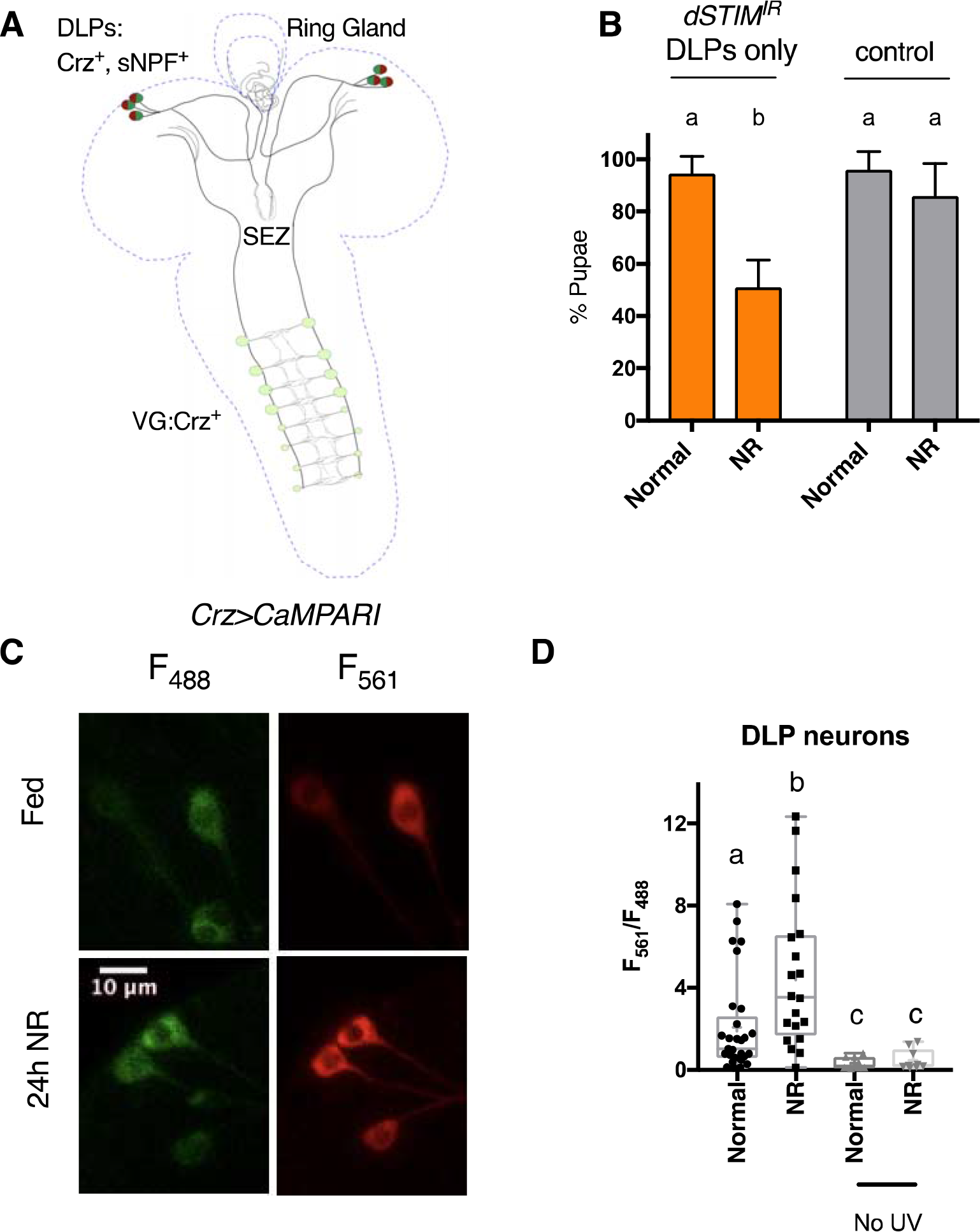
Crz^+^ and sNPF^+^ DLPs are required for development on NR media and activated by NR. **(A)** Cartoon of Crz^+^ and sNPF^+^ neurons in the larval CNS marked by *Crz-GAL4*. DLP: dorso lateral peptidergic; VG: Ventral Ganglion; SEZ: Subesophageal zone **(B)** % Pupae when *dSTIM* is selectively down-regulated only in Crz^+^ and sNPF^+^ DLPs, by using the *tsh-Gal8o* transgene and in the presence of *dicer2*. Control: *tshGal80*/+;*dSTIM*^*IR*^/+. Data represents mean ± SEM **(C)** Representative image. Expression of the UV-activated Ca^2+^ indicator, CaMPARI in Crz^+^ and sNPF^+^ DLPs, in larvae on 24 hours of normal (Fed) or NR media (24b NR). Fluorescence at 561m (F_561_) reflects Ca^2+^ levels, while at 488nm (F_488_) reflects levels of the indicator CaMPARI. **(D)** Quantification of Ca^2+^ levels as reported by F_561/488_ ratio in DLPs in larvae on 24 hours of normal or NR media, in the presence and absence of UV-stimulation. N>7 larvae for UV-stimulated; N=3 for No UV stimulation. Bars with the same alphabet represent statistically indistinguishable groups. Two-way ANOVA with Sidak multi comparison test p<0.05 for (B). Mann-Whitney Test for (D). See also Figure Supplement 2.

### *dSTIM* regulates NP synthesis and release in Crz neurons

Crz peptide levels were measured in DLP neurons by staining larval brains with an antiserum raised against the mature Crz peptide sequence [33]. Two locations on the DLP neurons were chosen for measurement: neuronal cell body/soma and neurite projections on the RG. In control DLP neurons, 24 hrs of NR caused average levels of Crz levels to increase, in both locations (Fig. 3A, S3A,B). In comparison, DLP neurons expressing *dSTIM*^*IR*^ displayed increased Crz peptide levels on normal food itself, and this remained unaltered upon NR, for both locations (Fig. 3A, S3B, C). sNPF levels could not be similarly measured by immunofluorescence because sNPF is expressed in many neurons close to the DLPs (Fig. S1C), making measurements specifically from the DLP soma difficult to quantify. Instead, semi-quantitative, direct, mass spectrometric profiling of dissected RGs was employed. This technique can measure peptide levels relative to stable isotopic standards at single neurohaemal release sites [34]. As Crz levels between the cell bodies and projections correlated, and Crz^+^ DLPs are the sole contributors of sNPF on the RG [24], this technique allowed us to infer sNPF levels in DLPs. In controls, 24hrs of NR, increased the average level of sNPF ~5-fold on the RG (Fig. 3C). In comparison, RG preparations from larvae where DLPs express *dSTIM*^*IR*^, displayed increased sNPF levels on normal food itself, and this remained unaltered upon NR (Fig. 3C). Although Crz was detected in the RG preparations, it was of much lower intensity. Average Crz levels increased with NR in the control, and in *dSTIM*^*IR*^ condition, but statistically higher levels of Crz were seen only in the NR, *dSTIM*^*IR*^ condition (Fig. S3E). Nonetheless, broad agreement in trends, between Crz using immunofluorescence and sNPF using MALDI-MS, suggest that the two peptides are similarly regulated by NR and *dSTIM*. This is consistent with genetic experiments which showed that over-expression of *dSTIM* can rescue loss of both, sNPF as well as Crz (Fig. 1C,D).

Thus, increased activation of DLP neurons by NR (Fig. 2D), appears to result in peptide accumulation. Loss of *dSTIM* increases peptide levels on normal food, and prevents an increase in peptide levels upon NR.

As an ER-Ca^2+^ sensor, *dSTIM* may potentially regulate several cellular processes that would affect NPs such as their synthesis, processing, trafficking and/or release. As *STIM*^*IR*^ increased peptide levels in the cell body as well as neurite projections on the RG, a major trafficking defect was unlikely (Fig. 3A vs S3B,C). This does not rule out a role for *dSTIM* in dense-core vesicle trafficking, but merely indicates that trafficking of Crz is not observably disrupted by *STIM*^*IR*^. We therefore proceeded to examine systemic and cellular phenotypes when molecules known to reduce overall NP synthesis (*InR*^*IR*^; Insulin Receptor) [35], peptide processing (*amon*^*IR*^) [25], and vesicle exocytosis (*Ral*^*DN*^) [17] were expressed in Crz neurons. All three perturbations caused a pupariation defect on NR media (Fig. 3D). However, despite similar systemic outcomes, *amon*^*IR*^, which reduces the prohormone convertase required for peptide maturation, reduced Crz levels (Fig. 3E). Because this is not seen with *STIM*^*IR*^, a role for dSTIM in peptide processing was not pursued further.

Expression of *InR*^*IR*^ caused a modest increase in Crz peptide levels in DLP neurons in the fed state and peptide levels did not increase as in the control, in NR media (Fig. 3F). InR is a global protein synthesis regulator, and its expression scales DIMM^+^ NE cell size, with functional consequences [35]. As the Crz^+^ DLPs are DIMM^+^ [36], we expected *InR*^*IR*^ to reduce, not increase peptide levels. A potential explanation for this observation is that Crz peptide levels are under feedback regulation, which was substantiated when we examined how Crz transcript and peptide levels are connected. Similarities between *InR*^*IR*^ and *STIM*^*IR*^ phenotypes, coupled with a previous observation that IP_3_R, another SOCE component, positively regulates protein synthesis in *Drosophila* neuroendocrine cells [22], prompted us to test if *dSTIM* too regulates protein synthesis in general. We ectopically expressed a physiologically irrelevant neuropeptide construct (ANF∷GFP), that yields a processed peptide in *Drosophila* neurons [31]. ANF∷GFP levels in control and *dSTIM*^*IR*^ DLP neurons were similar (Fig. S3E), suggesting that dSTIM does not have generic effects on peptide synthesis. Instead, its effect on Crz and sNPF synthesis may be specific. The lack of Crz elevation upon NR, in DLP neurons where *InR*^*IR*^ is expressed, leads to the speculation that InR signalling is required for the up-regulation of protein synthesis needed for increased peptide synthesis, processing and packaging during NR.

We previously found that *Ral* expression lies downstream of dSTIM-mediated SOCE in *Drosophila* pupal brains [37], and in dopaminergic neurons, over-expression of *Ral*^*DN*^ reduces secretion of ANF∷GFP [17]. These previous data, coupled with the observation that *Ral*^*DN*^ and *dSTIM*^*IR*^ show similarly high Crz levels in the fed state, suggest that *dSTIM* affects vesicle secretion through regulation of *Ral* expression. To independently validate if vesicle release is important in Crz neurons, we over-expressed a temperature sensitive dynamin mutant (*Shibire*^*ts*^) also shown to reduce NP release [38] and tetanus toxin (*TNT*) shown to prevent release of eclosion hormone, a neuropeptide [39]. Both manipulations caused a pupariation defect on NR (Fig. S3F). It is unclear why *Ral*^*DN*^ causes Crz levels to decrease upon NR. In *Drosophila* pacemaker neurons, *Ral* has been shown to bias the sensitivity of a neuropeptide receptor, the Pigment Dispensing Factor Receptor [40]. Perhaps, functions distinct from *Ral*’s contribution to vesicle exocytosis contribute to this observation. Nonetheless, the lack of increase in Crz levels in DLP neurons upon NR, when *InR*^*IR*^ and *Ral*^*DN*^ are expressed, assumes significance in the context of pupariation of NR larvae. Control larvae subject to 24hrs of NR, display increased Crz (Fig. 3A) and sNPF (Fig. 3C) levels, and then proceed to successfully complete development to pupae (Fig. 1C). Whereas, in d*STIM*^*IR*^, *InR*^*IR*^ and *Ral*^*DN*^ conditions, on NR, neither do DLPs display increased Crz levels (Fig. 3A, 3F), nor do all larvae pupariate (Fig. 1B, 1C, 3D). Thus, an increase in peptide levels on NR correlates with larval ability to pupariate on NR. Taken in context with increased neuronal activation during NR (Fig. 2D), and evidence that functional vesicle exocytosis (Fig. 3D, S3F) as well as adequate peptide production (Fig. 3D) is required for survival on NR, these data suggest that increased production and release of Crz and sNPF during NR, is required for NR larvae to successfully complete development.

To prove that Crz and sNPF are released during NR, the ideal experiment would be to measure levels of secreted NPs. But small size (8-10 amino acids), low hemolymph titres and high complexity of hemolymph, make peptide measurements by biochemical means highly challenging in *Drosophila*. Moreover, NPs exhibit endocrine as well as paracrine signalling [41]; and the latter will not be reflected in hemolymph measurements. Fortunately, in the Crz signalling system there is feedback compensation between secreted Crz and Corazonin receptor (*CrzR*) mRNA levels, providing a means to gauge secreted Crz levels indirectly. In adults, expression of *Crz*^*IR*^ in Crz neurons, increased levels of *CrzR* on the fat body [42]. We thus tested if *CrzR*, which in larvae appears to be expressed in the salivary glands and CNS [43], was subject to similar feedback. In larval brains, reducing Crz, using two different *Crz*^*IR*^ strains, not only caused a reduction in *Crz* mRNA levels (Fig. 3G), but also a concomitant increase in *CrzR* mRNA levels (Fig. 3H). Conversely, reducing CrzR levels (*CrzR*^*IR*^) in the larval CNS, results in up-regulation of *Crz* mRNA (Fig. S3G). This confirmed the existence of feedback in the Crz signalling system, and the use of neuronal *CrzR* transcript levels as a measure of secreted Crz levels. In line with this inference, we observed an up-regulation of *CrzR* mRNA in larval brains where Crz neurons are expressing either *InR*^*IR*^ or *amon*^*IR*^ or *Ral*^*DN*^ (Fig. 3I). Therefore, the observation that in the *STIM*^*IR*^ condition *CrzR* mRNA levels are high (Fig. 3I), supports the idea that dSTIM function is necessary for the secretion of optimal levels of Crz.

Because dSTIM-mediated SOCE is known to induce changes in gene expression [37], we probed if *Crz* expression is sensitive to NR and *dSTIM*. In the control, NR did not change *Crz* mRNA levels (Fig. 3J), suggesting that a post-transcriptional mechanism is responsible for increasing Crz peptide levels upon NR (Fig. 3A,S3B,C). In the *STIM*^*IR*^ condition, *Crz* transcript levels were up-regulated on normal food conditions (Fed) and no further increase was observed upon NR (Fig. 3J). The straightforward explanation for high Crz peptide levels in *STIM*^*IR*^ condition could therefore be attributed to higher gene expression of *Crz*. However, data from other perturbations in Crz neurons suggested that a linear interpretation between *Crz* mRNA and peptide levels cannot not be made. When *Crz*^*IR*^ is expressed, *Crz* mRNA is reduced (Fig. 3G), but peptide levels are elevated (Fig. S3I); whereas, when *amon*^*IR*^ is expressed, *Crz* mRNA is increased (Fig. S3H), but peptide levels are decreased (Fig. 3E). Meanwhile, in three conditions, *Crz* mRNA as well as peptide levels are higher than controls: *CrzR*^*IR*^(Fig. S3G vs S3I), *InR*^*IR*^(Fig. S3H vs Fig. 3F) and *Ral*^*DN*^ (Fig. S3H vs Fig 3F). Note also that both higher (*dSTIM*^*IR*^, *InR*^*IR*^, *Ral*^*DN*^) or lower (*amon*^*IR*^) Crz peptide levels in DLP cell bodies, result in reduced systemic Crz signalling (*CrzR* mRNA levels; Fig. 3I). These data indicate that Crz transcription, translation and release are independently regulated. A simple explanation for elevated levels of *Crz* transcript as well as peptide levels in *STIM*^*IR*^ is therefore, feedback compensation. Moreover, there is no change in Crz mRNA upon 24hrs of NR, when *STIM*^*IR*^ is expressed (Fig. 3J). Together, this argues against a direct role for *dSTIM* in regulating *Crz* gene expression.

In summation, these data have been inferred as follows: on normal food, partial loss of *dSTIM* reduces systemic Crz signalling, indicating a requirement for *dSTIM* in Crz secretion. On NR media, Crz^+^ DLPs are stimulated to increase peptide synthesis and release, in order for NR larvae to complete development. Peptide up-regulation upon NR is abrogated when *dSTIM* is reduced. These add up to suggest that *dSTIM* compromises NE cell function in a manner that affects peptide synthesis and release, with functional consequences for survival on NR.

**Fig 3.**
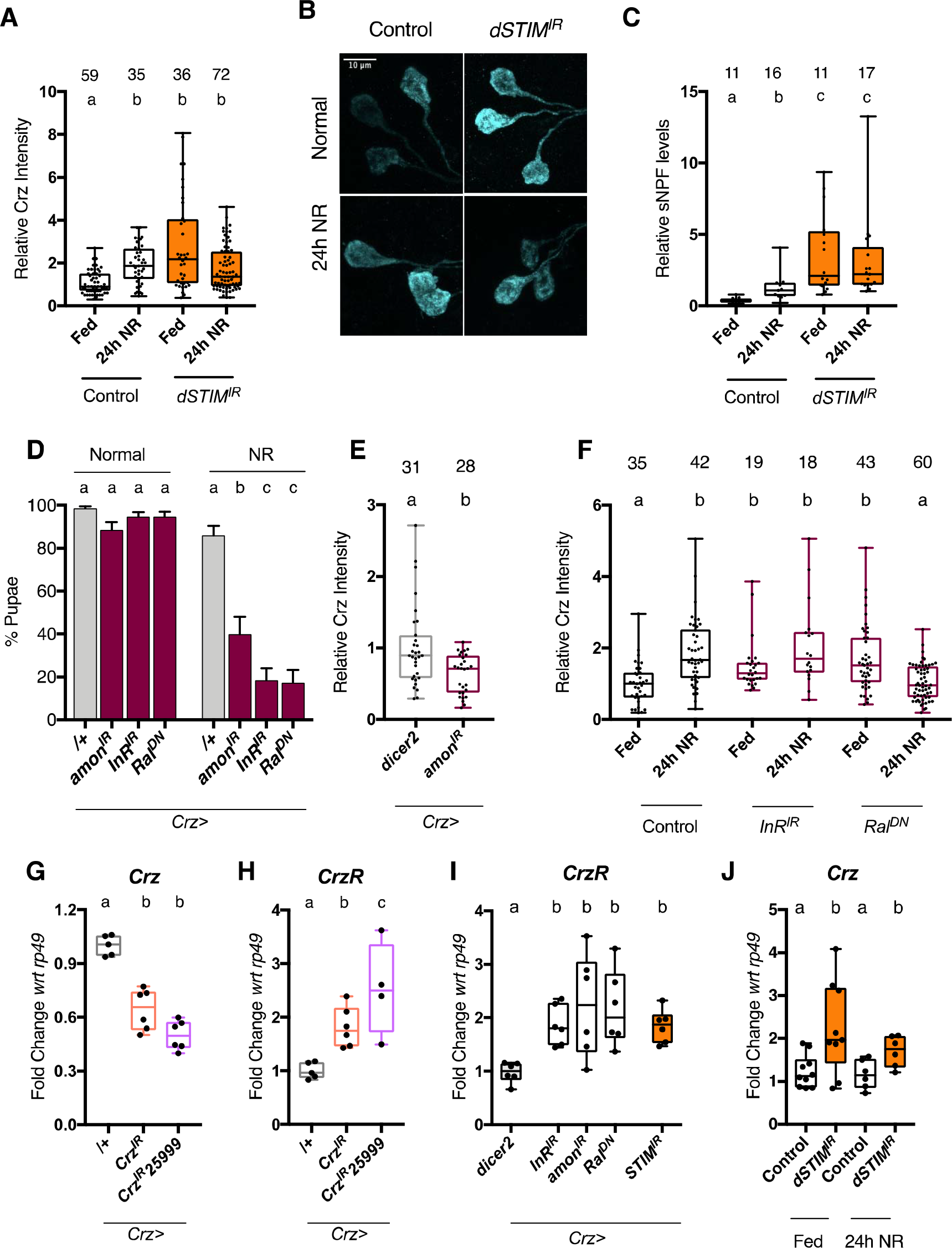
*dSTIM* regulates Crz and sNPF levels. Larvae were subjected to 24 hours of normal (fed) or nutrient restricted (NR) media. Crz levels were measured in DLP neurons by immunofluorescence on larval brains. All manipulations were performed using the *Crz-GAL4* driver **(A)** Relative levels of Crz peptide in DLP neuron cell bodies, Control=*crz*>*dicer2*. *dSTIM*^*IR*^=crz> *dSTIM*^*IR*^,*dicer2*. Number of cells measured shown atop bars. N>12 brains **(B)** Representative images for cell bodies measured in (A). **(C)** Relative levels of total sNPF peptides measured on dissected ring glands (N atop bars) and quantified using MALDI-MS. Externally added heavy standard (Hug-PK*) was used to normalise peptide levels between samples. **(D)** % Pupae on normal or NR media, upon reduced peptide processing (*amon*^*IR*^,*dicer2*) protein synthesis (Insulin receptor; *InR*^*IR*^) or vesicle exocytosis (dominant-negative *Ral*; *Ral*^*DN*^) in Crz^+^ neurons. Data represents mean ± SEM **(E)** Relative levels of Crz upon expression of *amon*^*IR*^ and *dicer2*. N>10 brains. **(F)** Relative levels of Crz upon indicated cellular perturbation of Crz^+^ neurons. N≥6 brains, control: *Crz-GAL4*/+. **(G)** *Crz* mRNA levels from larval brains when *Crz* is reduced by two different RNAi lines. N ≥ 5. **(H)** Corazonin receptor (*CrzR*) mRNA levels from larval brains with reduced *Crz*. N ≥ 4. **(I)** *CrzR* mRNA levels from larval brains expressing indicated cellular perturbations in Crz neurons. N ≥ 6 **(J)** *Crz* mRNA levels from larval brains. Control=*crz*>*dicer2*. *dSTIM*^*IR*^=*crz*> *dSTIM*^*IR*^,*dicer2* N ≥ 6. Bars with the same alphabet represent statistically indistinguishable groups. Kruskal-Wallis Test with Dunn’s multicomparison correction p<0.05 for (A), (C), (F). Mann-Whitney Test for(E). Two-way ANOVA with Sidak’s multi comparison test p<0.05 for (D), (J). One-way ANOVA with Tukey multi comparison test p<0.05 for(G), (H), (I). See also Figure Supplement 3.

### Systemic and cellular phenotypes observed with reduced *dSTIM* in Crz neurons can be rescued by increasing synthesis and release of peptides

To validate a role for dSTIM in peptide synthesis and release, we tested genetic perturbations that can compensate for this deficiency, to rescue developmental and cellular phenotypes associated with *dSTIM*^*IR*^ expression in Crz neurons. In the case of NPs, genetic over-expression may not translate to enhanced release, as proteins involved in NP processing as well as the regulated secretory pathway would need to be up-regulated. Furthermore, regulatory feedback from peptides to their transcription may complicate over-expression, as seen for the Crz signalling system (Fig. S3I). To get around these issues, and because *InR*^*IR*^ phenocopied *STIM*^*IR*^ (Fig. 3A,D,F,I, S3H), we opted to increase protein synthesis by over-expression of the Insulin receptor (*InR*). Cell size (Fig. S4A) as well as Crz levels (Fig. S4B,C) in DLP neurons scaled with *InR* over-expression, supporting the effectiveness of *InR.* To increase release, we over-expressed *Ral*^*WT*^ as *Ral* over-expression can compensate for vesicle release in dopaminergic neurons expressing *STIM*^*IR*^ [17]. In Crz neurons with reduced *dSTIM*, over-expression of either *InR* or *Ral*^*WT*^ rescued pupariation on NR media (Fig. 4A); restored peptide up-regulation upon NR (Fig. 4A,B) and decreased *CrzR* mRNA back to control levels (Fig. 4D). Of note, over-expressing *Ral*^*WT*^ or *InR* by itself, in Crz neurons, did not alter *CrzR* mRNA levels (Fig. 4D), suggesting that neuronal activation, which happens on NR media (Fig. 2D) potentiates their activity.

To increase neuronal activity, we utilised the temperature and voltage-gated cation channel, TrpA1 [44]. Over-expression of *dTrpA1* and its activation by raising the temperature to 30°C for 24 hours, in the *dSTIM*^*IR*^ background, rescued pupariation of NR larvae (Fig. 4E). It also restored the ability of DLP neurons to increase Crz levels upon NR (Fig. 4F) and decreased levels of *CrzR* mRNA (Fig. 4G). In line with the feedback between *CrzR* mRNA levels and systemic Crz signaling, over-expression of *dTrpA1* alone in Crz neurons resulted in a decrease in *CrzR* mRNA levels (Fig. 4G), supporting the role for neuronal activation in secreting Crz. Interestingly, development to adulthood on NR, for *dSTIM*^*IR*^ larvae, was also significantly increased upon over-expression of *InR* (Fig. S4E) and *TrpA1* (Fig. S4F), but not Ral^WT^ (Fig. S4E).

An optogenetic approach, utilizing the over-expression of Channelrhodopsin (ChR2-XXL), a light-sensitive cation channel, also rescued pupariation but not to the same extent (Fig. S4D). Poorer rescue with *CHR2-XXL* could be because sustained activation of this channel depress synaptic transmission and the channel is less conductive for Ca^2+^ compared to TrpA1 [45]. Similar genetic manipulation, with TrpA1 and Chr2-XXL, in a hypomorphic IP_3_R mutant (*itpr*^*ku*^) resulted in a small but significant rescue of *itpr*^*ku*^ pupariation on NR media (Fig. S4G,H).

Together, these results strongly suggested that defects arising from dysregulated intracellular Ca^2+^ signalling, may be overcome by increasing vesicle exocytosis (*Ral*^*WT*^, *TrpA1, ChR2-XXL rescue*) or protein synthesis (*InR* rescue). Importantly, the rescues observed with *InR*, *Ral*^*WT*^ and *dTrpA1* are effective at the molecular (*CrzR* levels), cellular (Crz peptide levels upon NR) as well as systemic (NR larvae) level.

**Fig 4.**
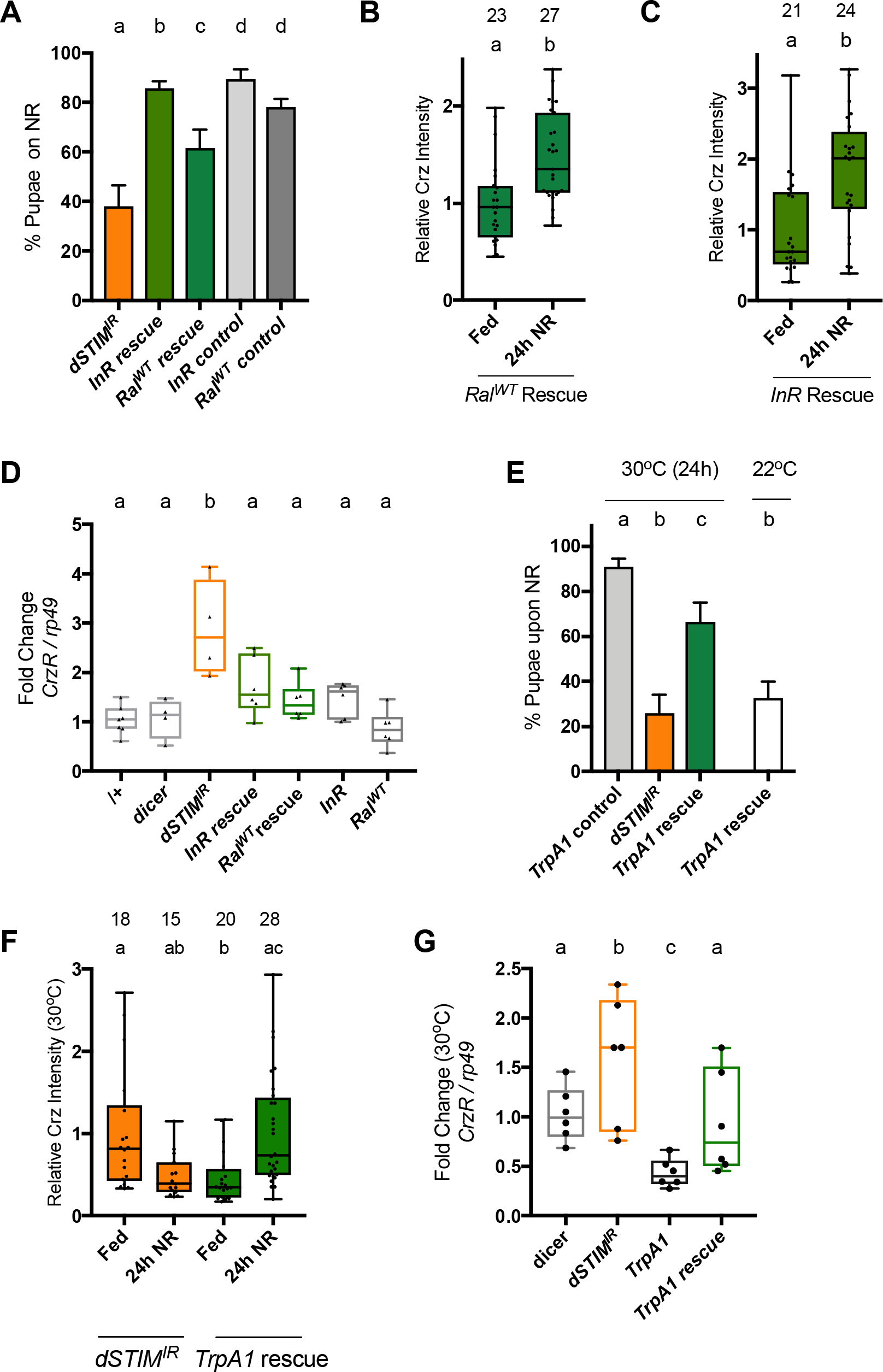
Systemic and cellular phenotypes caused by partial loss of dSTIM in Crz neurons, can be rescued by increasing peptide synthesis or release. **(A)** % Pupae upon expression of Insulin receptor (*InR*) or Ral (*Ral^WT^*) in Crz neurons expressing *dSTIM*^*IR*^. Genotypes: *dSTIM*^*IR*^ = *crz*>*dSTIM*^*IR*^. InR *rescue*=*Crz*>*InR*,*dSTIM*^*IR*^,*dicer2*. *Ral*^*WT*^rescue=*Crz*> *Ral*^*WT*^, *dSTIM*^*IR*^,*dicer2*. *InR* control: *dicer2;InRI*+; *dSTIM*^*IR*^ /+;. *Ral^IR^* control: *Ral^WT^*/+; *dSTIM*^*IR*^ /+. Data represents mean ± SEM. **(B)** *Ral^WT^*rescue or **(C)** InR rescue larvae were transferred to normal (Fed) or NR media for 24 hours. Crz immunofluorescence levels in DLP neurons were measured. Number of cells measured shown atop bars. N>7 brains **(D)** *Crz* receptor *(CrzR)* mRNA levels in larval brains expressing various molecules in Crz neurons. InR=*Crz*>*InR*. *Ral*^*WT*^ =*Crz*>*Ral*^*WT*^. N≥4. **(E)** % Pupae upon expression of TrpA1 in Crz neurons expressing *dSTIM^IR^*. TrpA1 control: *dicer2;TrpA1*/+, *dSTIM*^*IR*^ /+; TrpA1 rescue: *Crz*>*TrpA1*, *dSTIM*^*IR*^,*dicer2*. Larvae were reared at 25°C, at 88h±3h AEL age transferred to NR media and incubated either at 30°C or 22°C for 24 hours, and returned to 25°C thereafter. Data represents mean ± SEM. **(F)** Crz levels in DLP neurons upon 24 hrs of NR or normal food (Fed) at 30°C for indicated genotypes. N > 5 brains. **(G)** *Crz* receptor *(CrzR)* transcript levels in larval brains expressing various molecules in Crz^+^ neurons, when larvae are reared at 30°C. TrpA1: *Crz*>*TrpA1*. N=6. one-way ANOVA with a post hoc Tukey’s test p<0.05 for (A), (D), (E), (G). Mann-Whitney test for (B), (C). Kruskal-Wallis Test with Dunn’s multicomparison correction p<0.05 for (F). See also Figure Supplement 4.

## Discussion

This study employed an *in vivo* approach coupled to a functional outcome, in order to broaden our understanding of how STIM regulates neuropeptides. A role for dSTIM-mediated SOCE in *Drosophila* neuroendocrine cells for survival on NR was previously established [22]. The previous study offered the opportunity to identify SOCE-regulated peptides, produced in these neuroendocrine cells, that could be investigated in a physiologically relevant context.

In *Drosophila*, both Crz and sNPF have previously been attributed roles in many different behaviours. Crz has roles in adult metabolism and stress responses [42,46–48], sperm transfer and copulation [49], and regulation of ethanol sedation [50,51]. While, sNPF has been implicated in various processes including insulin regulation (Kapan et al., 2012; Lee et al., 2008) circadian behaviour [53], sleeping [54,55] and feeding [27]. Thus, the identification of Crz and sNPF in coping with nutritional stress is perhaps not surprising, but a role for them in coordinating the larval to pupal transition under NR is novel.

A role for Crz in conveying nutritional status information was originally proposed by Jan Veenstra [56], which this study now supports. In larvae, Crz^+^ DLPs are known to play a role in sugar sensing [57] and in adults, they express the fructose receptor Gr43a [58]. Additionally, they express receptors for neuropeptides DH31 [59], DH44 [59] and AstA [56], which are made in the gut as well as larval CNS. Together, these observations and our study are strongly indicative of a role for Crz^+^ DLPs in directly or indirectly sensing nutrients, with a functional role in larval survival and development in nutrient restricted conditions.

Several neuropeptides and their associated signalling systems are evolutionarily conserved [11,12]. The similarities between Crz and GnRH (gonadotrophin-releasing hormone), and sNPF and PrRP (Prolactin-releasing peptide), at the structural [11], developmental [60] and receptor level therefore, is intriguing. Structural similarity of course does not imply functional conservation, but notably, like sNPF, PrRP has roles in stress response and appetite regulation [61]. This leads to the conjecture that GnRH and PrRP might play a role in mammalian development during nutrient restriction.

dSTIM regulates Crz and sNPF at the levels of peptide release and likely, peptide synthesis upon NR. We speculate that neuroendocrine cells can use these functions of STIM, to fine tune the amount and timing of peptide release, especially under chronic stimulation (such as 24hrs NR), which requires peptide release over a longer timeframe. Temporal regulation of peptide release by dSTIM may also be important in neuroendocrine cells that co-express peptides with multifunctional roles, as is the case for Crz and sNPF. It is conceivable that such different functional outcomes may require distinct bouts of NP release, varying from fast quantile release to slow secretion [62]. As elevation in cytosolic Ca^2+^drives NP vesicle release, neurons utilise various combinations of Ca^2+^ influx mechanisms to tune NP release. For example, in *Drosophila* neuromuscular junction, octopamine elicits NP release by a combination of cAMP signalling and ER-store Ca^2+^, and the release is independent of activity-dependent Ca^2+^ influx [63]. In the mammalian dorsal root ganglion, VGCC activation causes a fast and complete release of NP vesicles, while activation of TRPV1 causes a pulsed and prolonged release [64]. dSTIM-mediated SOCE adds to the repertoire of mechanisms that can regulate cytosolic Ca^2+^ levels and therefore, vesicle release. This has already been shown for *Drosophila* dopaminergic neurons [17] and this study extends the scope of release to peptides. Notably, dSTIM regulates exocytosis via Ral in neuroendocrine cells, like in dopaminergic neurons.

In *Drosophila* larval Crz^+^ DLPs, dSTIM appears to have a role in both fed, as well as NR conditions. On normal food, not only do Crz^+^ DLPs exhibit small but significant levels of neuronal activity (Fig. 2D) but also, loss of dSTIM in these neurons reduced Crz signalling (Fig. 3I). Thus, dSTIM regulates Ca^2+^ dynamics and therefore, neuroendocrine activity, under basal as well as stimulated conditions. This is consistent with observations that basal SOCE contributes to spinogenesis, ER-Ca^2+^ dynamics as well as transcription [65]. However, in our case, this regulation appears to have functional significance only in NR conditions as pupariation of larvae, with reduced levels of *dSTIM* in Crz^+^ neurons, is not affected on normal food (Fig. 1B). In a broader context, STIM is a critical regulator of cellular Ca^2+^ homeostasis as well as SOCE, and a role for it in the hypothalamus has been poorly explored. Because STIM is highly conserved across the metazoan phyla, our study predicts a role for STIM and STIM-mediated SOCE in peptidergic neurons of the hypothalamus. There is growing evidence that SOCE is dysregulated in neurodegenerative diseases [66], In neurons derived from mouse models of familial Alzheimer’s’ disease [67] and early onset Parkinson’s [65], reduced SOCE has been reported. How genetic mutations responsible for these diseases manifest in neuroendocrine cells is unclear. If they were to also reduce SOCE in peptidergic neurons, it’s possible that physiological and behavioural symptoms associated with these diseases, may in part stem from compromised SOCE-mediated NP synthesis and release.

## Material and Methods

### Fly Husbandry

Flies were grown at 25°C in 12h:12h L:D cycle. Normal food: **(1L recipe: 80g corn flour, 20g Glucose, 40g Sugar, 15g Yeast Extract, 4mL propionic acid, *p*-hydroxybenzoic acid methyl ester in ethanol 5mL, 5mL ortho butyric acid)** For nutritional stress assay, flies were allowed to lay eggs for 6 hours on normal food. After 88hours, larvae were collected and transferred to either normal or NR (100mM Sucrose) food. Pupae and adults were scored after 10 days of observation.

### Fly strains

*Canton S* was used as the wild type (+) control.

The following strains were obtained from Bloomington Drosophila Stock Centre: *AKH-GAL4* (25684), *Crz-GAL4* (51976), *DSK-GAL4* (51981), *sNPF-GAL4* (51981), *UAS-Crz^IR^*25999 (25999), *UAS-dicer2* (24651), *UAS-TeTxLc* (28837), *UAS-TeTxLc-IMP* (28838), *UAS-CaMPARI* (58761), *UAS-GFP^nls^* (4776), *UAS-mRFP* (32218), *UAS-TrpA1* (26263), *UAS-InR* (8248), *UAS-amon^IR^* (28583), *UAS-Ral*^*DN*^ (32094)

The following strains were obtained from Drosophila Genetic Resource Center, Kyoto: *sNPF-GAL4* (113901)

The following were from Vienna Drosophila Research Centre stock collection: *UAS-IP*_3_*R*^*IR*^ (106982), *UAS-STIM*^*IR*^ (47073), *UAS-Crz^IR^* (30670), *UAS-InR^IR^* (999)

The following was from Exelixis at Harvard Medical School: *sNPF*^*00448*^ (C00448)

The following were kind gifts: *AstA*^*1*^-*GAL4* (David Anderson), *dILP2-GAL4* (Eric Rulifson), *hug-GAL*4 (Michael Pankratz), *NPF-GAL4* (Ping Shen), *UAS-sNPR*^*IR*^ (K weon Yu), Crz∷mCherry (Gábor Juhász), *UAS-hid∷UAS-rpr* (Tina Mukherjee), *UAS-Shibire^ts^* (Toshihiro Kitamoto), *tsh-GAL80* (Julie Simpson), *UAS-preproANF∷GFP* (Edwin Levitan), *UAS-CHR2XXL* (Robert Kittel and Georg Nagel)

The following were previously generated in our laboratory: *itpr*^*ka1091*^, *itpr*^*ug3*^, *UAS-Orai*^*E180A*^, *UAS-itpr*^+^,*UAS-Stim*, *UAS-Ral*^*WT*^

### Larval Feeding

Ten 3^rd^ instar larvae were collected and placed on cotton wool soaked with solution of 4.5% dissolved yeast granules and 0.5% Erioglaucine (Sigma, 861146). Controls contained no dye. Feeding was allowed for 2 hours at 25°C. 5 larvae per tube were crushed in 100μL of double distilled water. Solution was spun at 14000 rpm for 15 minutes and 50μL was withdrawn for absorbance measurement at 625nm in a 96-well plate. 5μL was used to measure protein content using the Pierce BCA Protein Assay kit (#23227).

### qRT-PCR

RNA was isolated from 12-15 larval brains at the specified time points using Trizol. cDNA synthesis was carried out as described [26]. All mRNA levels are reported as fold change normalized to *rp49*. Primer sequences:

*rp49*, F:CGGATCGATATGCTAAGCTGT, R:GCGCTTGTTCGATCCGTA.
*Crz*, F:TCCTTTAACGCCGCATCTCC, R:CGTTGGAGCTGCGATAGACA
*CrzR*, F:CTGTGCATCCTGTTTGGCAC, R:GGCCTTGTGTATCAGCCTCT

### Measuring neuronal activation using *CaMPARI*

Early third instar larvae were transferred to either normal or NR food. After 24 hours, larvae were recovered and immobilized on double sided tape. UV light from a Hg-arc lamp was focused using the UV filter, on the larvae through a 10X objective on Olympus BX60, for 2 minutes. Larvae were them immediately dissected in ice-cold PBS, mounted in PBS and imaged using Olympus FV-3000 Confocal microscope using a 40X objective and high-sensitivity detectors. Microscope settings for laser intensity, PMT settings and magnification were kept identical for all measurements. Each experiment always had a no UV control, in which larvae were subject to immobilisation but not UV light. Fluorescence intensity was calculated for each cell body using Image J.

### Immunofluorescent staining

For expression patterns, 3rd instar larval brains with RGs attached were dissected in ice-cold PBS and fixed in 3.7% formaldehyde at 4°C for 20mins. The samples were washed 4 times in PBS and mounted in 60% glycerol. Endogenous fluorescence was acquired on Olympus FV-3000 using a 20X, 40X or 60X objective, and processed used ImageJ. For samples requiring antibody staining brains were similarly processed and then subjected to permeabilisation (0.3% Triton X-100 + PBS; PBSTx) for 15 mins, 4 hr blocking in 5% normal goat serum in PBSTx at 40C, followed by overnight incubation in primary antibody (1:1000 Chicken-GFP, Abcam: ab13970) and secondary with Alexa 488 or Alexa 594 (1:400; Abcam). For corazonin (1:1000; raised in Rabbit; Jan Veenstra, University of Bordeaux), all the above steps remained the step, except that dissected brains were fixed for 1hr at RT in 4% PFA and the secondary was anti-rabbit Alexa 405 (1:300, Abcam). Cell bodies were outlined manually and integrated density was used to calculate CTCF (Corrected Total Cell Fluorescence). For all samples, a similar area was measured for background fluorescence.

### Direct peptide-profiling by MALDI-TOF MS

Ring glands were dissected in cold HL3.1 and transferred to a MALDI plate as previously described [68], 0.2 μl of matrix (saturated solution of recrystallized α-cyano-4-hydroxycinnamic acid in MeOH/EtOH/water 30/30/40% v/v/v) was added, containing 10 nM of stable isotope-labeled HUG-pyrokinin (HUG-PK* (Ser–Val[d8]–Pro–Phe–Lys–Pro–Arg–Leu–amide, Mw = 950.1 Da; Biosyntan, Berlin, Germany)) and 10 nM labeled myosuppressin (MS* (Thr–Asp–Val[d8]–Asp–His–Val–Phe–Leu–Arg–Phe–amide, Mw = 1255.4 Da; Biosyntan) MALDI-TOF mass spectra were acquired in positive ion mode on a 4800 Plus MALDI TOF/TOF analyzer (MDS Sciex, Framingham, MA, USA) in a mass range of 900-4000 Da and fixed laser intensity with 20 subspectra and 1000 shots per sample. Data were analyzed with Data Explorer 4.10. Spectra were baseline corrected and de-isotoped. The sum of the resulting relative intensities of the de-isotoped peaks was calculated for the different ion adducts (H^+^, Na^+^, K^+^) of each peptide as well as the labeled peptides*. Then, the ratios sNPF/HUG-PK* and corazonin/MS* were calculated, using the labeled peptide with the most similar molecular weight. For sNPF, all isoforms (1/2-short, 1-long, -3 and -4) variants were totaled.

### Optogenetic and thermogenic experiments

For thermogenetic (*dTrpA1*, *Shibire*^*ts*^) experiments, larvae were matured to 88hours AEL at 25°C. After transfer to either NR or normal food, vials were placed at 22°C, 25°C or 30°C for either 24 hours (*dTrpA1*) or till the end of observation time (*Shibire*^*ts*^). For optogenetic experiments (*Chr2-XXL*), larvae were matured to 88AEL in the dark. After transfer to either NR or normal food, one set was placed in the dark while another was placed in an incubator with regular white lights that were on continuously till the end of observation time.

## Author contributions

M, C.W and G.H designed research; M performed research, except MALDI-MS which was performed by C.W.; M, C.W and G.H. analysed data; M wrote the paper with inputs from C.W. and G.H.

## Acknowledgments

Supported by Wellcome Trust/ DBT India Alliance (Early Career Award #IA/E/12/1/500742 to M) and NCBS core funding (GH). We thank Jan Veenstra for the Crz antibody and helpful suggestions with immunostaining. Members of the Hasan lab for help with fly transfers, critical comments and helpful discussions. Drosophila fly community (List in supplementary information), BDSC, VDRC, NIG and DGRC for fly strains. NCBS Central Imaging and Flow Facility, for imaging, and Jörg Kahnt (core facility for mass spectrometry and proteomics, Max-Planck-Institute for Terrestrial Microbiology, Marburg) for access to the mass spectrometer.

