## supplementary information for "“ER-Ca^2+^ sensor STIM regulates neuropeptides required for development under nutrient restriction in *Drosophila*”"

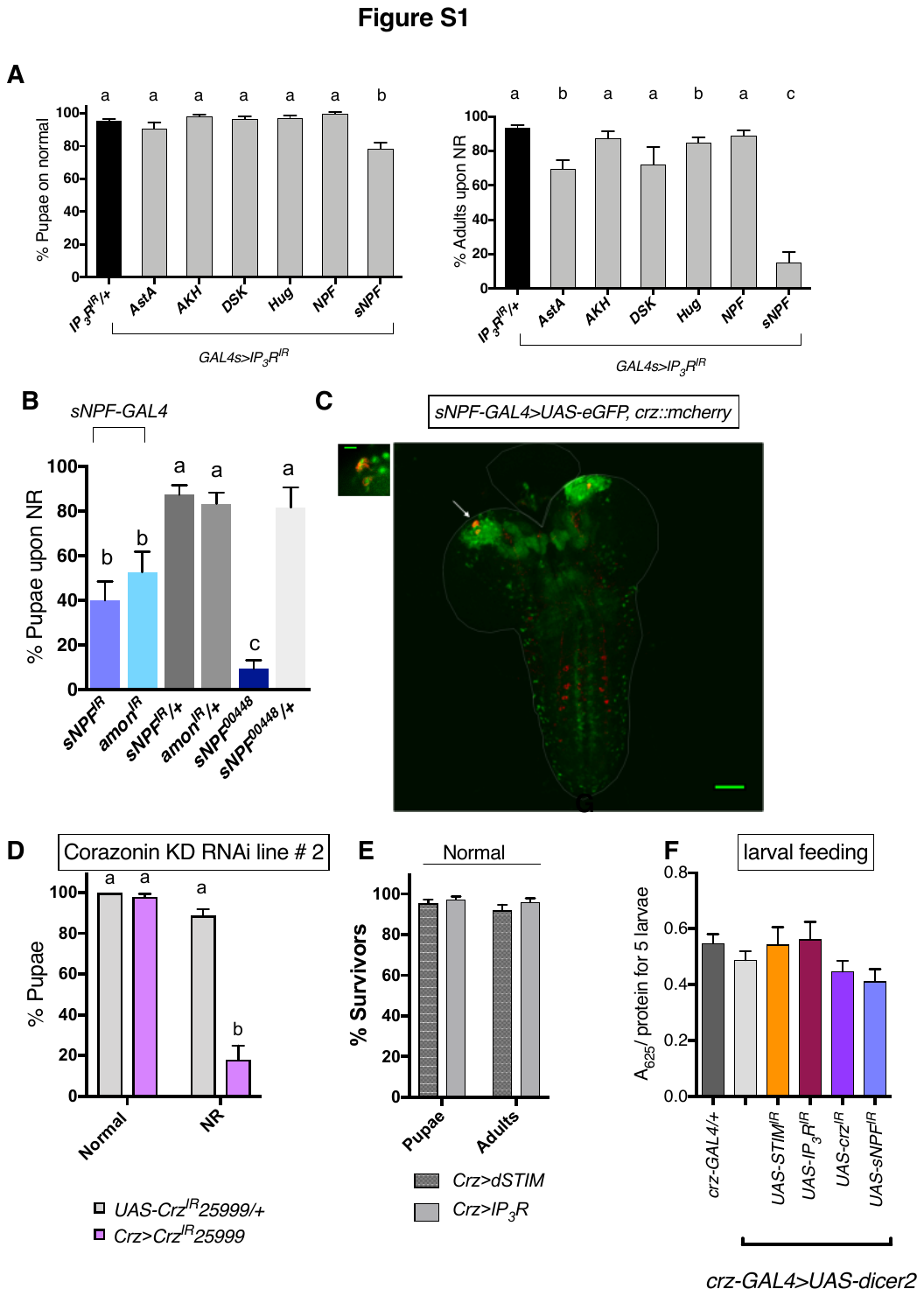

**Fig S1**

**A.** Genetic Screen. GAL4s for various peptides were used to drive *IP_3_R^IR^* and the corresponding larvae were tested for their ability to pupariate on normal or NR media. AstA: Allatostatin A; AKH: Adipokinetic Hormone; DSK: Drosulphakinin; Hug: Hugin; NPF: Neuropeptide F; sNPF: short Neuropeptide F.

**B.** % Pupae in NR media when sNPF is reduced by RNAi (*sNPR^IR^*) or by reducing an enzyme required for neuropeptide processing (*amon^IR^*) in *sNPF-GAL4* expressing cells, or in a hypomorphic sNPF mutant (*sNPF^00448^*).

**C.** 3^rd^ instar larval brain expressing GFP in *sNPF-GAL4* producing neurons and a mcherry-tagged corazonin (*Crz::mcherry*). Note the co-localisation of *sNPF-GAL4* with Crz::mcherry, only in the brain lobes (arrow).

**D.** % Pupae on normal and NR media when Crz is reduced in Crz^+^ neurons, using a second RNAi line (*crz^IR^25999*)

**E.** % Pupae and adults that developed on normal food from larvae where Crz^+^ neurons over-expressed either *dSTIM* or *IP_3_R*. Differences are not statistically significant.

**F.** Food intake as measured by absorbance of coloured food (A_625_) fed to larvae for the indicated genotypes. N= 8 sets of 5 larvae each. Differences are not statistically significant.

one-way ANOVA with a post hoc Tukey’s test p<0.05 for (A), (B), (F). Ordinary two-way ANOVA with Sidak’s multi-comparison test p<0.05 for (D). Bars with the same alphabet represent statistically indistinguishable groups. Data represents mean ± SEM**.**

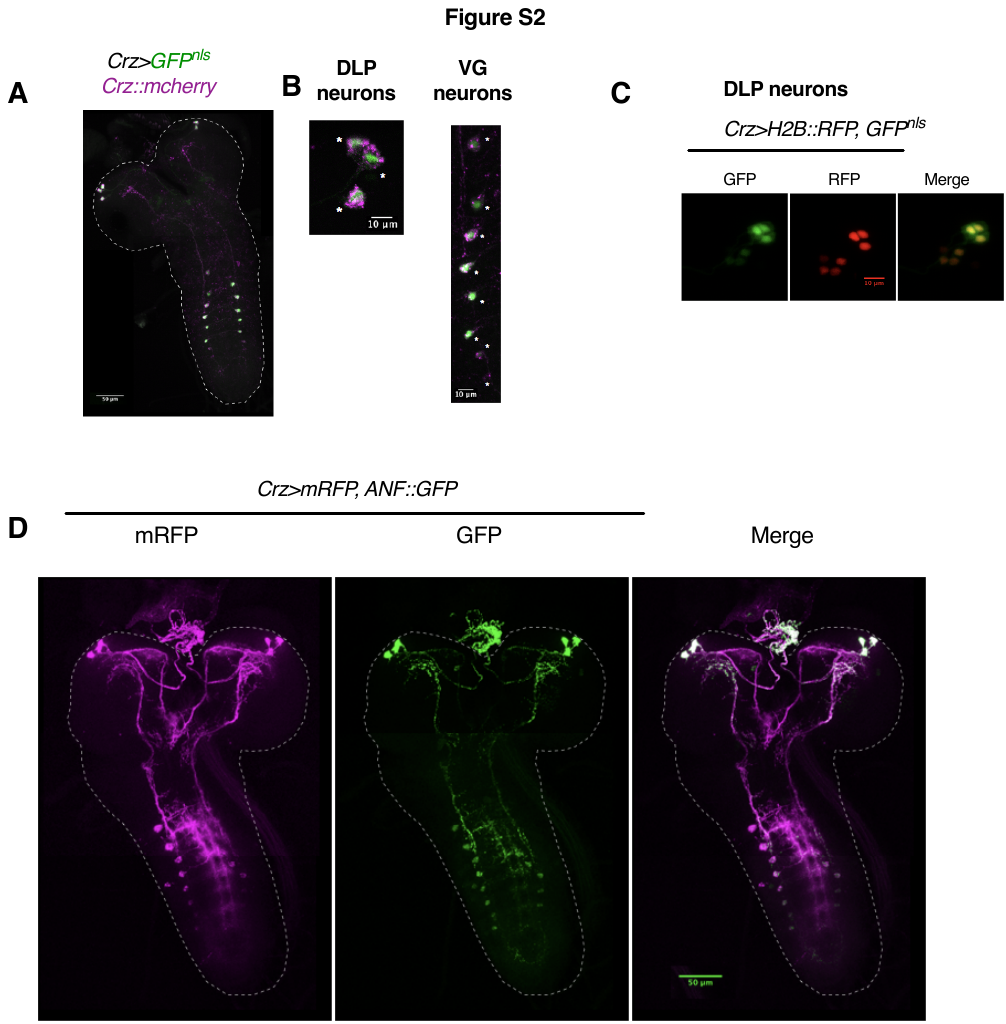

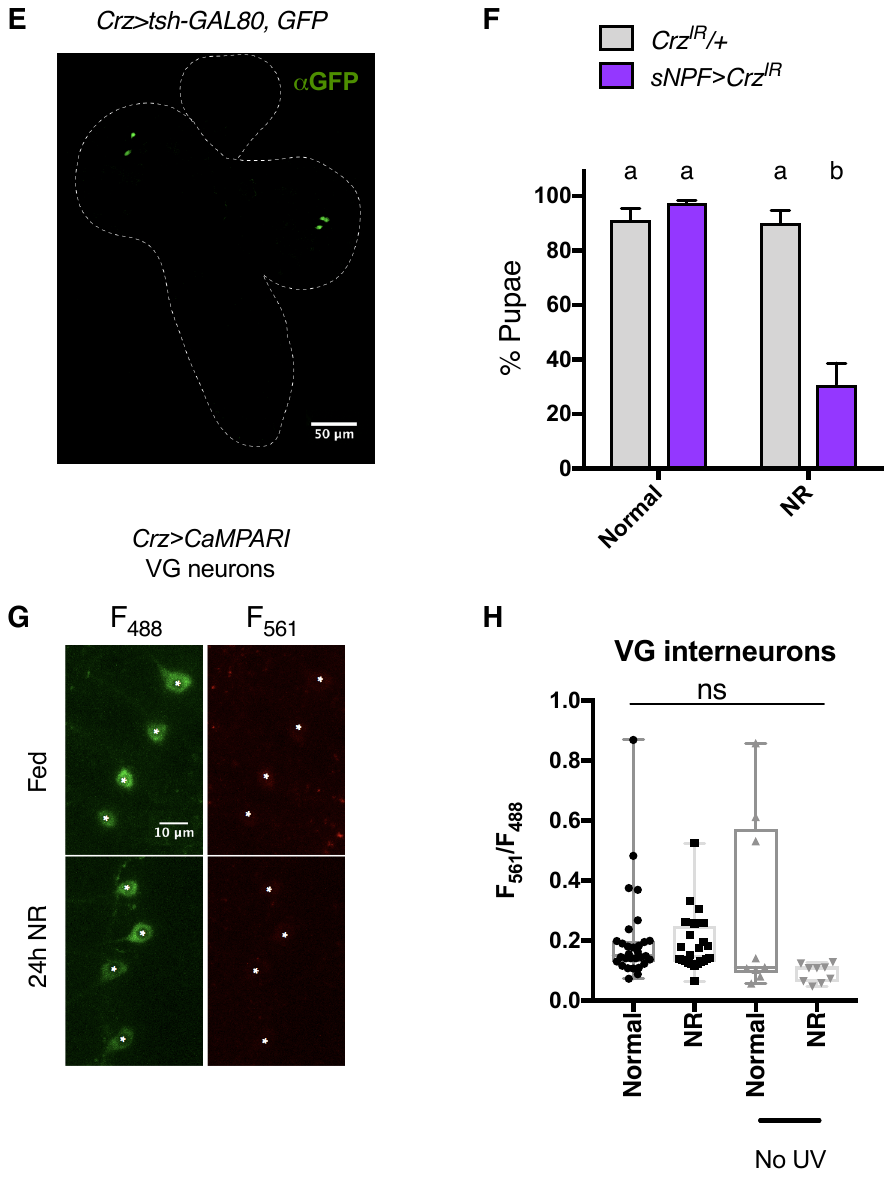

**Fig S2**

**A. and B.** Expression pattern of *Crz-GAL4* (observed via *GFP^nls^*) closely matched the expression of Crz (observed by a genomically integrated, mCherry-tagged corazonin: Crz::mCherry).

**C.** *Crz-GAL4* also expresses in ~4-5 additional neurons in the DLP region, that are not marked by Crz::mCherry (See A). The level of expression also varies with the marker. *GFP^nls^ vs* histone-tagged to RFP (H2B::RFP).

**D.** Overall distribution of NPs in Crz^+^ neurons followed by the expression of GFP-tagged rat ANF (ANF::GFP) and mCD8-tagged membrane bound RFP (*UAS*-*mRFP).* Note the exclusion of ANF from projections that end in the SEZ.

**E.** Representative image. Expression of *tsh-GAL80* in *crz>GFP* expressing brains causes the loss of GFP expression in the VG.

**F.** % Pupae when Corazonin is reduced (*Crz^IR^*) in *sNPF-GAL4* expressing neurons. This allows restricted expression of *Crz^IR^* only in DLPs (Fig. S1C). Ordinary two-way ANOVA with a post hoc Sidak’s multi-comparison test p<0.05. Data represents mean ± SEM.

**G**. Representative image. Expression of the UV-activated Ca^2+^ indicator, CaMPARI in Crz^+^ VG neurons. F_561_ reflects Ca^2+^ levels, while F_488_ reflects levels of the indicator.

**H.** Quantification of F_561/488_ ratio in the presence and absence of UV-stimulation, after 24hrs in either normal or NR food, in Crz^+^ VG neurons. N>7 larvae for UV-stimulated; N=3 for No UV stimulation. Kruskal-Wallis Test with Dunn’s multi-comparison correction p<0.05

Bars with the same alphabet represent statistically indistinguishable groups.

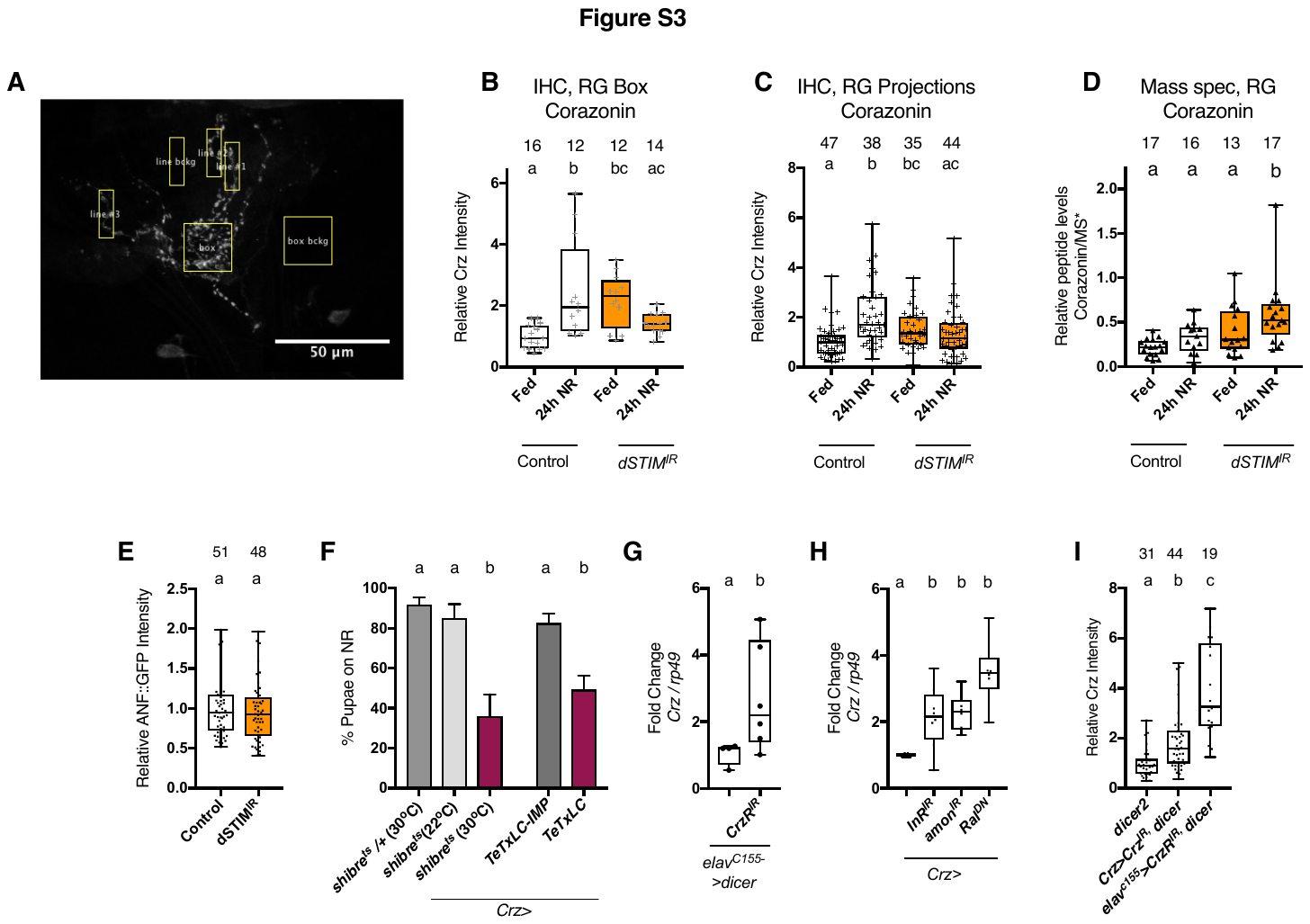
 **Fig S3**

**A.** Representative image. How “Box” and “projection” areas were delineated for measuring levels of Crz by immunofluorescence, at the ring gland (RG). Box is a 50X50 px square. Lines are a 50X15px rectangle. These measurements were made in a subset of DLPs for which Crz intensity were measured in cell bodies, plotted in Fig. 3A.

**B. and C**. Quantification of Crz levels at the RG.

**D.** Relative Crz peptide levels measured on dissected RGs (N atop bars) and quantified using MALDI-MS. Externally added heavy standard (MS*) was used to normalise peptide levels between samples. Kruskal-Wallis Test with Dunn’s multi-comparison correction p<0.05

**E.** Relative levels of the Rat Neuropeptide ANF::GFP measured by GFP fluorescence, in Crz^+^ DLPs in either control (*crz>dicer2*) or *dSTIM^IR^* condition (*crz>dstim^IR^,dicer2*)**.** Two-tailed t-test. Total number of cell bodies counted mentioned atop bars. N>15 brains.

**F.** % Pupae when vesicle release is perturbed either by expressing a dynamin mutant activated at 30^o^C (*Shibre^ts^),* or tetanus toxin light chain (TeTxLc). TeTxLC-IMP: inactivated TeTxLc. Ordinary one-way ANOVA with a post hoc Tukey’s test p<0.05. Data represents mean ± SEM.

**G**. *Crz* mRNA levels in larval brain with pan-neuronal reduction in *CrzR* (*crzR^IR^*). Two-tailed t-test. N=4.

**H.** *Crz* mRNA levels in larval brains where protein synthesis (*InR^IR^*) or peptide processing (*amon^IR^*) or vesicle exocytosis (*Ral^DN^*) are perturbed in Crz^+^ neurons. Ordinary one-way ANOVA with a post hoc Tukey’s test p<0.05. N>4.

**I**. Relative levels of Crz measured on cell bodies of Crz^+^ DLP neurons when either Crz (*Crz^IR^*) is reduced in Crz^+^ neurons or its receptor CrzR (*crzR^IR^*) is reduced pan-neuronally. Kruskal-Wallis Test with Dunn’s multicomparison correction p<0.05. N>10 brains.

Bars with the same alphabet represent statistically indistinguishable groups.

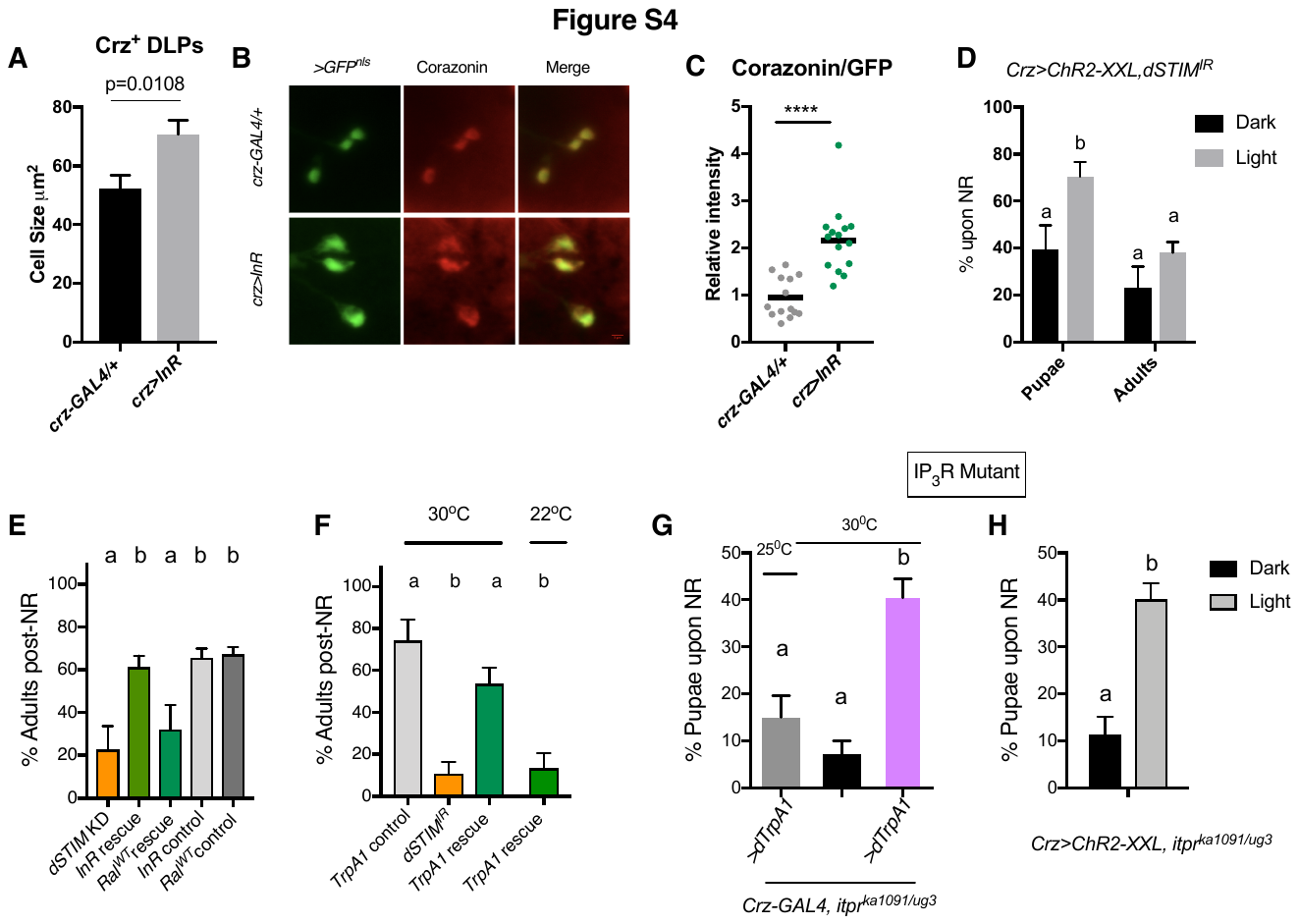
 **Fig. S4**

**A.** Cell size of DLP neurons, upon over-expression of Insulin receptor (*InR)* in Crz^+^ neurons. N = 5 brains. Student’s t-test. Representative images in **B**.

**C**. Relative Crz peptide levels in cell bodies of Crz^+^ DLP neurons expressing *InR*. N =5. Student’s t-test. p<0.001

**D.** % Pupae upon NR, when in Crz^+^ neurons, *dSTIM* was reduced, and *ChR2-XXL*, the light activated channel was ectopically expressed. Larvae were reared in the dark and post-transfer to NR, either continued to be kept in the “Dark” or moved to an incubator with 24 lights (white light) ON (Light) till the end of the pupariation assay (~ 10days). two-way ANOVA with a post hoc Sidak’s multi-comparison test p<0.05.

**E**. % Adults recovered upon over-expression of Insulin receptor (*InR*) or Ral (*Ral^WT^*) in Crz^+^ neurons expressing *dSTIM^IR^*. InR rescue: *crz>InR,dSTIM^IR^,dicer2. Ral^WT^* rescue: *crz> Ral^WT^, dSTIM^IR^,dicer2*. *InR* control: *dicer2;InR*/+; *dSTIM^IR^* /+;. *Ral^WT^* control: *Ral^WT^* /+; *dSTIM^IR^* /+. Ordinary one-way ANOVA with a post hoc Tukey’s test p<0.05.

**F**. % Adults recovered upon over-expression of TrpA1 in Crz^+^ neurons expressing *dSTIM^IR^*. TrpA1 control: *dicer2;TrpA1*/+; *dSTIM^IR^* /+; TrpA1 rescue: *crz>TrpA1, dSTIM^IR^,dicer2* Ordinary one-way ANOVA with a post hoc Tukey’s test p<0.05. N=6

Ordinary one-way ANOVA with a post hoc Tukey’s test p<0.05. N= 6.

**G and H**. % Pupae upon NR, in an hypomorphic IP_3_R mutant (*itpr^ka1091/ug3^*) with Crz^+^

neurons over-expressing either *dTrpA1*, and post-transfer incubating at 30^o^C for 24

hours, or *ChR2-XXL* and reared in light till the end of the assay. Ordinary one-way ANOVA with a post hoc Tukey’s test p<0.05. for (G). Student’s t-test for (H).

Bars with the same alphabet represent statistically indistinguishable groups. Data represents mean ± SEM.
